# Development of a SYBR Green quantitative PCR assay for detection of *Lates calcarifer herpesvirus* (LCHV) in farmed barramundi

**DOI:** 10.1101/2020.04.23.057018

**Authors:** Watcharachai Meemetta, Jose A. Domingos, Ha Thanh Dong, Saengchan Senapin

## Abstract

*Lates calcarifer* herpes virus (LCHV) is a new virus of farmed barramundi in Southeast Asia. However, a rapid detection method is yet to be available for LCHV. This study, therefore, aimed to develop a rapid quantitative PCR (qPCR) detection method for LCHV and made it timely available to public for disease diagnostics and surveillance in barramundi farming countries. A newly designed primer set targeting a 93-bp fragment of the LCHV putative major envelope protein encoding gene (*MEP*) was used for developing and optimizing a SYBR Green based qPCR assay. The established protocol could detect as low as 10 viral copies per µl of DNA template in a reaction containing spiked host DNA. No cross-amplification with genomic DNA extracted from host as well as common aquatic pathogens (12 bacteria and 3 viruses) were observed. Validation test of the method with clinical samples revealed that the virus was detected in multiple organs of the clinically sick fish but not in the healthy fish. We thus recommend that barramundi farming countries should promptly initiate active surveillance for LCHV in order to understand their circulation for preventing possibly negative impact to the industry.

**Highlights:**

- This study reported a new SYBR Green qPCR method for detection of LCHV
- The qPCR method had detection limit of 10 copies per µl plasmid DNA template when spiked with genomic DNA from the host
- The aforementioned method is highly specific to LCHV
- Validation with clinical samples revealed that LCHV could be detected from multiple organs with fin and brain the best organs for qPCR detection

## INTRODUCTION

Barramundi (*Lates calcarifer*) or Asian sea bass is one of the economically important finfish species in Asia-Pacific which has been farmed in a wide range of salinity in either open cage systems or earthen ponds (Jerry et al., 2014). Barramundi, like other intensively farmed fish, is susceptible to various infectious pathogens and often subject to serious outbreaks and economic losses (Dong et al., 2017a, b; Jerry et al., 2014; Ransangan et al., 2010; Toranzo et al., 2005). In recent years, three newly emerging viruses have been reported in farmed barramundi in Asia-Pacific, including scale drop disease virus (SDDV) (Gibson-Kueh et al., 2012; de Groof et al., 2015), *Lates calcarifer* herpes virus (LCHV) (Chang et al., 2017) and *Lates calcarifer* birnavirus (LCBV) (Chen et al., 2019). Both SDDV and LCHV were discovered from disease outbreaks where the fish showed clinical symptoms of “scale drop” and laboratory infections with the cultivated virus from cell culture resulted in up to 60% and 77% cumulative mortality, respectively (de Groof et al., 2015; Chang et al., 2017). By contrast, LCBV did not induce mortality in the controlled laboratory trial (Chen et al., 2019).

LCHV discovered by Chang et al. (2017) is a novel member of the family *Alloherpesviridae*, which is genetically most similar to *Ictalurid herpesvirus* 1 (<60% nucleotide identity), a pathogenic virus of channel catfish. LCHV is an enveloped virus with diameter of approximately 100 nm, and genome size of ∼130 kb while other members of *Alloherpesviridae* are between 150-250 nm in diameter and 100-250 kb in genome size (Hanson et al., 2011; Chang et al., 2017).

Both SDDV and LCHV infections cause similar scale drop disease-like gross signs which are clinically indifferentiable. Therefore, molecular detection methods are required for diagnostic and screening purposes. Several DNA-based detection methods for SDDV have been freely available such as single PCR (Senapin et al., 2019), semi-nested PCR (Charoenwai et al., 2019), loop-mediated isothermal amplification (LAMP) (Dangtip et al., 2019), probe-based qPCR (de Groof et al., 2015), and SYBR Green-based qPCR (Sriisan et al., 2020). The latter one is the most sensitive method with a detection limit of 2 copies of DNA template per reaction. In case of LCHV, following discovery of the virus, several primer sets for detection purpose were published in a patent (Chang et al., 2017), the use of these methods thus might be conditionally limited. According to requests from private sector, this study, therefore, developed a new, sensitive qPCR detection method for rapid diagnostics of LCHV and made it available to promote active surveillance for preventing wide-spread of this pathogen.

## MATERIALS AND METHODS

### Fish samples and DNA extraction

In 2019, there were 3 batches of barramundi samples subjected to testing for LCHV in our laboratory. Batch 1 comprised of adult fish (n = 5) in which 4 of them exhibited scale drop clinical signs while one fish had healthy looking appearance. Eight different tissue types (liver, kidney, spleen, gills, fin, brain, eyes and muscle) from each fish were dissected and individually preserved in 95% ethanol. Batch 2 comprised of apparently healthy barramundi fry that were ethanol-preserved. Three whole fry were pooled and considered as one sample for the test (n = 5 pools). Batch 3 (n = 10) comprised of ethanol-preserved spleen samples collected from 5 apparently healthy juvenile fish and 5 clinically sick fish showing scale drop disease-like symptoms. Approximately 5 mg tissue was subjected to DNA extraction using conventional sodium dodecyl sulfate/proteinase K containing lysis solution followed by phenol/chloroform extraction and ethanol precipitation. The obtained DNA pellet was resuspended in sterile distilled water and quantified using spectrophotometry at OD 260 and 280 nm.

### Primer design and PCR conditions

LCHV primers were designed to target a 93 bp partial fragment of a putative major envelop protein (*MEP*) gene of the virus. Forward primer LCHV-MEP93-qF: 5’-GTACTTCATCGCCTACGGAGC-3’ and reverse primer LCHV-MEP93-qR: 5’-TACGTGTGCTTGAGGAGGTC-3’ were synthesized from Bio Basic, Canada. Gradient PCR was firstly conducted to find an optimal annealing temperature (Ta) using Ta ranging from 58 to 65 °C. The reaction mixture of 20 µL contained 200 ng of DNA extracted from fin of LCHV-infected fish, 1x iTaq Universal SYBR Green SuperMix (Bio-Rad Cat.no. 172-5121) and 200 nM of each primer. The PCR amplification conditions were initial denaturation at 95°C for 10 min followed by 40 cycles of 95°C for 10 s and annealing at 58-65 °C for 30 s (Bio-Rad CFX Connect Real-Time PCR) followed by melt peak analysis. Finally, Ta of 61°C was selected and the same thermocycling conditions were used throughout the study.

### Sensitivity, qPCR efficiency, and specificity assays

Positive control plasmid namely pMEP93 was constructed for use in diagnostic sensitivity test. This was done by cloning the 93-bp *MEP* amplified fragment obtained above into pGEM T-easy vector (Promega) and transforming into *Escherichia coli* XL-1 blue. After colony PCR verification of potential correct clones, one recombinant clone was sent for DNA sequencing at Macrogen (South Korea). Copy number of pMEP93 was calculated based on plasmid size and concentration at https://cels.uri.edu/gsc/cndna.html and the pMEP93 was 10 fold-serially diluted from 10^7^ to 1 copies/µl. Plasmid dilutions (2 µl) were then used as template in qPCR conditions described above. To mimic a real test, each reaction also contained spiked 100 ng DNA extracted from a healthy barramundi. Control reaction without pMEP93 was used as a negative control. Analytical sensitivity experiment was conducted in 3 replicates within the same run. Standard curve was then automatically generated from quantification cycle (Cq) values being plotted versus log_10_ pMEP93 quantity. Formula for copy number calculation, coefficient of correlation (R^2^), and amplification efficiency (E) values were also provided by the Bio-Rad Maestro Software.

The optimized qPCR protocol was subsequently used to test for specificity against extracted genomic DNA from i) clinically healthy fish, ii) from 12 common aquatic bacterial species, and iii) from fish samples infected with either infectious spleen and kidney necrosis virus (ISKNV), nervous necrosis virus (NNV), or scale drop disease virus (SDDV). Sample sources and preparation were previously described (Charoenwai et al., 2019; Sriisan et al., 2020). DNA extracted from fin of LCHV-infected fish was used as positive control. No template control was used as negative reaction. Specificity test was performed in 2 replicates by 2 qPCR runs.

### LCHV detection in field samples

The newly developed qPCR was used to detect and quantify LCHV loads in the barramundi DNA samples prepared from 3 fish batches. 200 ng DNA template was used in each qPCR reaction. The obtained Cq was used to calculate viral copy numbers in the samples using the equation, copy number = 10^(Cq - Intercept)/Slope^ i.e. 10^(Cq – 41.34))/-3.539^ derived from the stand curve described above. Comparative evaluation of the viral loads in different fish tissue types was performed using samples from batch 1.

## RESULTS

### SYBR Green based LCHV qPCR

The LCHV qPCR protocol developed in this study had a detection limit of 10 copies/µl template i.e. 20 copies/reaction. Mean Cq ± SD values of the detection limit were 37.91 ± 0.33 (**Fig. 1a**). In other words, samples with Cq ≤ 37.91 ± 0.33 were considered as LCHV positive tests. The amplified products yielded uniform melting temperatures (Tm) at 84.0°C (**Fig. 1b**), indicating that the primers and the condition assayed were specific. The 93-bp amplicon had a relatively high Tm due to its 58% GC content of the sequence. Note that there was 1 nucleotide difference (**Supplemental Fig. 1**) between the target sequence in this study and that from the previous data (Chang et al., 2017). Based on the standard curve shown in **Fig. 1c**, the performance of the newly developed qPCR was high determined by its amplification efficiency (E) of 91.7% with R^2^ of 0.995. When evaluated the protocol specificity, the LCHV qPCR was demonstrated to be highly specific because it only detected LCHV infected sample but not DNA extracted from 3 other viruses, 12 bacteria, or clinically healthy fish tested. Data from one of the two replicates is shown in **Fig. 2**.

**Fig. 1.**
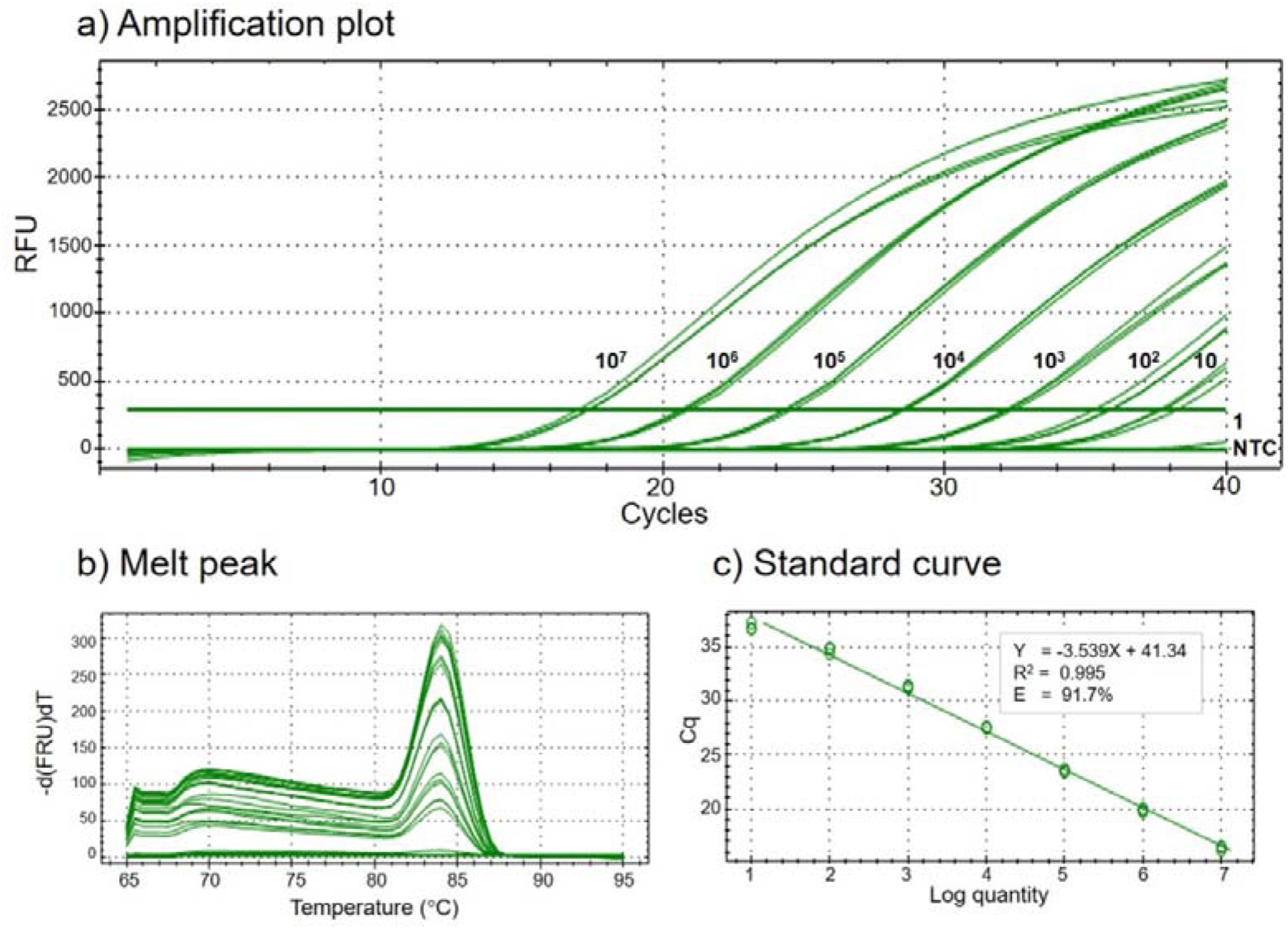
Performance and sensitivity of LCHV SBYR Green-based qPCR. (a) Amplification plots of positive control plasmid pMEP93 serial dilutions from 10^7^ to 1 copies containing 100 ng spiked fish DNA in each reaction. Three technical replicates were done for each dilution. (b) Melt peak analysis of the products obtained in (a). (c) Standard curve derived by plotting Cq values versus log_10_ pMEP93 concentrations. Formula for copy number calculation, R^2^ and E values are shown in the box.

**Fig. 2.**
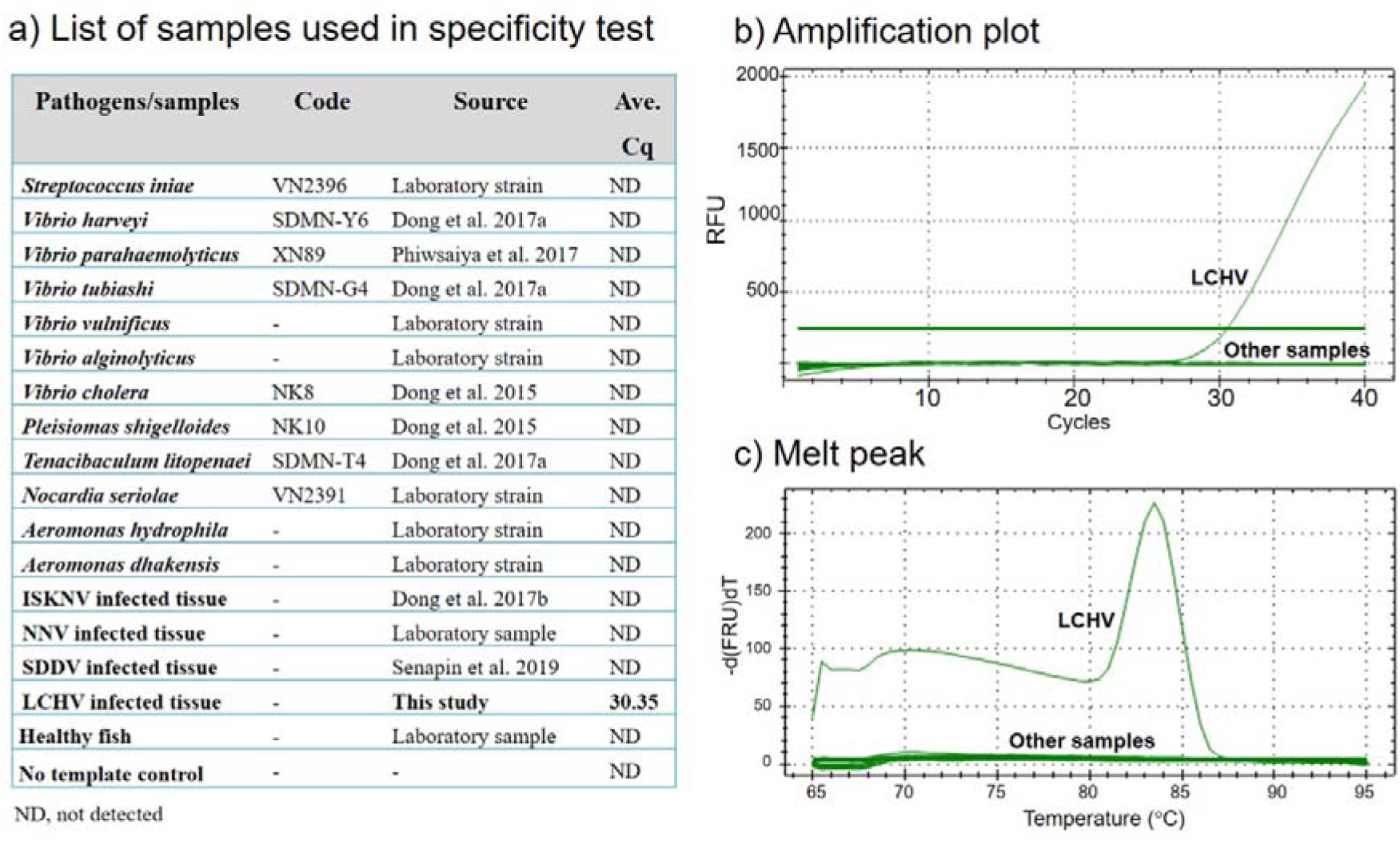
Specificity test of LCHV SBYR Green-based qPCR. (a) DNA samples extracted from bacterial isolates and viral infected fish as well as control reactions (DNA from healthy fish and no template control) were used in the specificity assay. Average Cq values from technical replicates are shown. ND, not detected. (b) Amplification plots and (c) melt peak analysis of products from samples shown in table (a).

### LCHV detection in fish samples

Tissue tropism of LCHV was revealed using the sample batch 1. Among all 8 tissues (liver, kidney, spleen, gills, fin, brain, eyes and muscle) tested from 4 diseased barramundi, the qPCR assay detected LCHV DNA at variable loads in 3-7 tissue types of each fish but not in the kidney samples (**Table 1**). There was only 1 in 4 liver sample which tested positive for LCHV with low viral loads (32 copies/200 ng DNA). DNA extracted from the fin, gills and muscle had averagely higher LCHV loads (24-597 copies/200 ng DNA) when compared to that of the brain, eyes, spleen and liver (13.7-184 copies/200 ng DNA). LCHV was not detected in any of the 8 tissue types of a clinically healthy fish from the same batch (**Table 1**).

**Table 1.** LCHV loads in 8 different tissues from 5 barramundi samples in batch 1.

| Sample | Fish clinical status | Cq and LCHV load*/200 ng template) |  |  |  |  |  |  |  |
| --- | --- | --- | --- | --- | --- | --- | --- | --- | --- |
|  |  | Liver | Kidney | Spleen | Gills | Fin | Brain | Eyes | Muscle |
| 1 | “Scale drop” | ND | ND | 33.33<br>[184] | 35.91<br>[34.3] | 34.35<br>[94.7] | 34.4<br>[82.4] | ND | ND |
| 2 | “Scale drop” | ND | ND | 36.05<br>[31.3] | 32.39<br>[339] | 34.26<br>[100] | ND | 35.07<br>[59.3] | 34.43<br>[89.9] |
| 3 | “Scale drop” | ND | ND | ND | ND | 33.65<br>[149] | 37.32<br>[13.7] | ND | 36.46<br>[24.0] |
| 4 | “Scale drop” | 36.02<br>[32] | ND | 35.55<br>[43] | 32.92<br>[240] | 31.52<br>[597] | 36.20<br>[28.4] | 34.81<br>[70.2] | 32.12<br>[404] |
| 5 | Healthy | ND | ND | ND | ND | ND | ND | ND | ND |

\*Cq values are the above number while the LCHV loads are shown in [ ]. Grey highlights
samples with LCHV load more than 90 copies. ND, not detected

The established qPCR was also applied to diagnose field samples from batches 2 and 3. DNA from all five pools of clinically healthy fry from batch 2 tested negative for by LCHV (**Table 2**). In batch 3, LCHV was detected from 5 clinically sick fish with viral loads ranging from 18.7 to 115.9 copies per 200 ng DNA (Cq 36.84-34.04) and undetectable in 5 clinically healthy fish (**Table 2**).

**Table 2.** LCHV detection test results of samples from batches 2 and 3.

| Batch no. | Sample no. | Fish clinical status | Tested tissue | Cq | LCHV loads/200 ng DNA template |
| --- | --- | --- | --- | --- | --- |
| 2<br>(fry) | Pool 1 | Healthy | whole body | ND | Negative test |
|  | Pool 2 | Healthy | whole body | ND | Negative test |
|  | Pool 3 | Healthy | whole body | ND | Negative test |
|  | Pool 4 | Healthy | whole body | ND | Negative test |
|  | Pool 5 | Healthy | whole body | ND | Negative test |
| 3<br>(juvenile) | Fish no. 1 | Healthy | spleen | ND | Negative test |
|  | Fish no. 2 | Healthy | spleen | ND | Negative test |
|  | Fish no. 3 | Healthy | spleen | ND | Negative test |
|  | Fish no. 4 | Healthy | spleen | ND | Negative test |
|  | Fish no. 5 | Healthy | spleen | ND | Negative test |
|  | Fish no. 6 | “Scale drop” | spleen | 35.39 | 48.2 |
|  | Fish no. 7 | “Scale drop” | spleen | 34.04 | 115.9 |
|  | Fish no. 8 | “Scale drop” | spleen | 35.48 | 45.4 |
|  | Fish no. 9 | “Scale drop” | spleen | 36.23 | 27.9 |
|  | Fish no. 10 | “Scale drop” | spleen | 36.84 | 18.7 |

ND, not detected

## DISCUSSION

LCHV and SDDV infections reportedly cause similar gross sign of “scale drop” in infected barramundi (de Groof et al., 2015; Chang et al., 2017). Despite the fact that both SDDV and LCHV have been recently discovered, the “scale drop” syndrome has been recognized in Southeast Asia since 1992 (Gibson-Kueh et al., 2012; de Groof et al., 2015). Therefore, it has raised a concern that both of these pathogens may have been long undiagnosed in farmed barramundi due to unavailability of respective diagnostic tools at that time. Nevertheless, currently several molecular detection methods for SDDV are available to support disease investigation. However, following discovery of LCHV as an emerging virus in Singapore in 2017 (Chang et al., 2017), there was no continuous research up-to-date. Although several sets of primers were described in the original patent document by Chang et al. (2017), their detection limits and test specificity remain uninvestigated. The validated qPCR method developed in this study might serve as a useful diagnostic tool for rapid screening of the suspected cases as well as active surveillance and early monitoring of the pathogen for the barramundi aquaculture industry.

Detection of LCHV in multiple organs of the clinically sick fish suggests that the virus caused systemic infection, similar to that of SDDV (Senapin et al., 2019; Charoenwai et al., 2019; Sriisan et al., 2020). Interestingly, the liver and kidney tissues which are normally used for PCR diagnostics of fish viruses appeared to be unsuitable for LCHV detection while the fin seemed to be the best targeted tissue due to its highest viral loads, followed by gills, muscle, spleen and brain. There was a limitation of fish numbers in this study, further comparative analysis should be done with larger sample numbers in order to gain a better understanding of virus tissue tropism as well as viral loads in the fish at different stages of infection. Nevertheless, this knowledge might be useful for establishment of cost-effective and non-destructive sampling strategies of fin and/or gills of farmed fish for periodical monitoring of the LCHV.

The present study focused primarily on the development and validation of a sensitive qPCR detection method for LCHV. Apart from LCHV, several pathogens have been reported to cause similar clinical signs of “scale drop” including SDDV (Gibson-Kueh et al., 2012; de Groof et al., 2015; Senapin et al., 2019), a pathogenic strain of *Vibrio harveyi*, and *Tenacibaculum maritimum* (Dong et al., 2017a; Gibson-Kueh et al., 2012). Relatively low viral loads present in the clinically sick fish with scale drop disease-like symptoms suggests that LCHV might be an opportunistic pathogen rather than the true causative agent of the diseased fish investigated in this study. However, identification of other pathogens in field samples was not done in this study. We thus recommend that investigation of the at least four aforementioned agents should be considered for the fish showing scale drop symptoms in order to weigh involvement of each pathogen in field outbreaks.

## Acknowledgements

This study was supported by a research grant from Mahidol University.

## Tables and Figures

**Supplemental Fig. 1.**
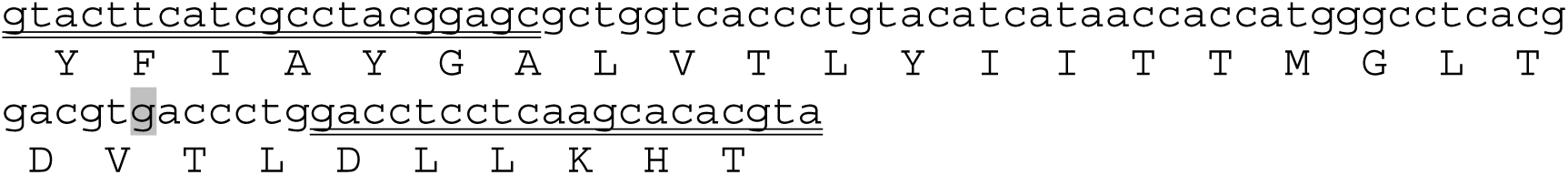
Nucleotide sequence of the LCHV qPCR target. Putative translated amino acid sequence is shown in capital alphabets. qPCR primers (double underlined) were designed to generate a 93-bp fragment of LCHV *MEP* gene. Compared to previously documented sequence (Chang et al. 2017), there is one silent mutation (gray highlighted) found in this study.

